# Membrane proteins with high N-glycosylation, high expression, and multiple interaction partners were preferred by mammalian viruses as receptors

**DOI:** 10.1101/271171

**Authors:** Zheng Zhang, Zhaozhong Zhu, Wenjun Chen, Zena Cai, Beibei Xu, Zhiying Tan, Aiping Wu, Xingyi Ge, Xinhong Guo, Zhongyang Tan, Zanxian Xia, Haizhen Zhu, Taijiao Jiang, Yousong Peng

**Affiliations:** College of Biology, Hunan University, Changsha, China; College of Computer Science and Electronic Engineering, Hunan University, Changsha, China; Center of System Medicine, Institute of Basic Medical Sciences, Chinese Academy of Medical Sciences & Peking Union Medical College, Beijing, China; Suzhou Institute of Systems Medicine, Suzhou, China; School of Life Sciences, Central South University, Changsha, China; State Key Laboratory of Chemo/Biosensing and Chemometrics, Hunan University, Changsha, China

## Abstract

Receptor mediated entry is the first step for viral infection. However, the relationship between viruses and receptors is still obscure. Here, by manually curating a high-quality database of 268 pairs of mammalian virus-host receptor interaction, which included 128 unique viral species or sub-species and 119 virus receptors, we found the viral receptors were structurally and functionally diverse, yet they had several common features when compared to other cell membrane proteins: more protein domains, higher level of N-glycosylation, higher ratio of self-interaction and more interaction partners, and higher expression in most tissues of the host. Additionally, the receptors used by the same virus tended to co-evolve. Further correlation analysis between viral receptors and the tissue and host specificity of the virus shows that the virus receptor similarity was a significant predictor for mammalian virus cross-species. This work could deepen our understanding towards the viral receptor selection and help evaluate the risk of viral zoonotic diseases.

## Introduction

In the new century, much progress has been made in prevention and control of infectious diseases, but the recent serial outbreaks of Zika virus ^[1]^, Ebola virus (EBOV) ^[2]^ and Middle East Respiratory Syndrome Coronavirus (MERS-CoV) ^[3]^ indicate that the viral infectious diseases still pose a serious threat to human health and global security. The virus is the most abundant biological entity on Earth and exists in all habitats of the world ^[4]^. Nearly all cellular organisms are prey to viral attack. Humans were reported to be infected by hundreds of viruses ^[5, 6]^. Most of the human emerging infectious diseases are zoonotic, with viruses that originate in mammals of particular concern ^[7]^, such as the Human Immunodeficiency Virus (HIV) ^[8]^ and Severe Acute Respiratory Syndrome Coronavirus (SARS-CoV) ^[9]^. Mammals are not only the most closely related animal to humans in phylogeny, but also contact with humans most frequently ^[7]^, especially for the livestock and pet. For effective control of human viral diseases, much attention should be paid to the mammalian virus.

Receptor-binding is the first step for viral infection of host cells ^[10-13]^. Multiple types of molecules could be used as viral receptors ^[12, 14]^, including protein ^[15-17]^, carbohydrate ^[18, 19]^ and lipid ^[20]^. How to select receptors by the virus is an important unsolved question ^[13, 14, 16, 21]^. Specificity and affinity are two most important factors for viral receptor selection ^[14]^. Carbohydrates and lipids are widely distributed on host cell surfaces and easy targets for viruses to grab ^[10, 11]^. Compared to these molecules, proteins were reported to be more suitable receptors because of stronger affinity and higher specificity for viral attachment, which could increase the efficiency of viral entry and facilitate viruses to expand their host ranges and alter their tropisms ^[10-12^, ^14, 15]^ Previous studies have shown that proteins that were abundant in the host cell surface or had relatively low affinity for their natural ligands, were preferred by viruses as receptors, such as proteins involved in cell adhesion and recognition by reversible, multivalent avidity-determined interactions ^[10, 15]^. This suggests that the selection of proteins by viruses as receptors should not be a random process. A systematic analysis of the characteristics of the viral receptor could help understand the mechanisms under the receptor selection by viruses.

The virus-receptor interaction was reported to be a principal determinant of viral host range, tissue tropism and cross-species infection ^[11, 16, 22]^. The existence and expression of the virus receptor in a host (or tissue) should be a prerequisite for viral infection of the host (or tissue) ^[21]^. Usually, a virus mainly infects some particular type of hosts or tissues. For example, the influenza virus mostly infects cells of the respiratory tract ^[23]^. However, the virus-receptor interaction is a highly dynamic process. Some viruses can recognize one or more receptors ^[13, 14, 24]^, which can also differ among virus variants or during the course of infections ^[14, 25, 26]^. In some cases, a few amino acid mutations in the viral protein or the receptor could abolish or enhance viral infection ^[27-29]^. Besides, the virus-receptor interaction is under continuous evolutionary pressure to increase the viral infection efficiency, which may result in the emergence of virus variants with altered host or tissue tropism. For example, the SARS-CoV and MERS-CoV, which belong to the same genus, *betacoronavirus*, have evolved to use different receptors (angiotensin I converting enzyme 2 (ACE2) and dipeptidyl peptidase 4 (DPP4) respectively) and also infect different hosts ^[11, 16, 28]^. Despite of numerous studies about the tissue and host specificity of the virus and the viral receptor, the systematical correlation characteristics between them are still obscure.

Here, by manually curating a high-quality database of 268 pairs of mammalian virus-receptor interaction, which included 128 unique viral species or sub-species and 119 virus receptors, we systematically analyzed the structural, functional, evolutionary and tissue-specific expression characteristics of mammalian virus receptors, which could not only deepen our understanding towards the mechanism behind the viral receptor selection, but also help to predict and identify viral receptors. Besides, we also investigated the associations between the tissue and host specificity of the virus and the viral receptor, and further evaluated the risk of viral cross-species based on viral receptors. It would help for early warning and prediction of viral zoonotic diseases.

## Results

### Database of mammalian virus-host receptor interaction

To understand how the virus selects receptors, we manually curated a high-quality database of 268 pairs of mammalian virus-host receptor interactions (Figure 1 and Table S1), which included 128 unique viral species or sub-species from 21 viral families and 119 virus receptors from 13 mammal species. The viral receptor collected here belonged to 13 mammal species (Figure S1A), among which the human accounted for the most (74/119). The viruses included in the database covered all groups of viruses in the Baltimore classification (Figure 1). Among them, the single-stranded RNA (ssRNA) virus accounted for over half of all viruses (76/128), while the double-stranded RNA (dsRNA) virus accounted for the least (3/128). On the level of family, the family of *Picornaviridae* of ssRNA virus, *Retroviridae* of Retro-transcribing viruses (RT) and *Herpesviridae* of double-stranded DNA (dsDNA) viruses were the most abundant ones in the database (Figure 1 and Table S1).

**Figure 1.**
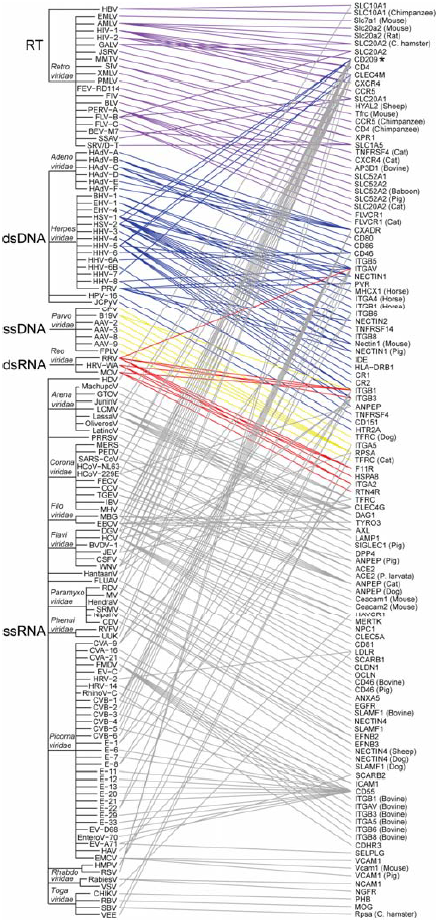
The mammalian viruses and their related receptors in our database. The lines between the virus and their related receptors were colored according to the group of the virus in the Baltimore classification. The names of some viral families were presented in italic. Viral names were displayed in abbreviation (see Table S1 for the full name). The host names were given for the receptor of non-human mammal species. The receptor CD209 was marked with an asterisk. For more details about the mammalian virus and their receptors, please see the website http://www.computationalbiology.cn:5000/viralReceptor.

### Association between mammalian viruses and their receptors

Analysis of the association between the virus and their receptors showed that 60% of viruses (77/128) used only one receptor (Figure 1 and Figure S1B), while the remaining viruses used two or more receptors. Surprisingly, some viruses, such as the Human alphaherpesvirus 1 (HSV-1) and Hepacivirus C (HCV), used more than five receptors. We next analyzed the receptor usage on the level of viral family. For fifteen viral families including two or more viruses in the database, all of them used two or more sets of receptors, suggesting that different viruses of the same family tend to use different receptors. For example, in the family of *Togaviridae*, the Chikungunya virus (CHIKV), the Rubella virus (RBV) and the Sindbis virus (SBV) used the receptor of prohibitin (PHB), myelin oligodendrocyte glycoprotein (MOG) and ribosomal protein SA (RPSA), respectively. On the other hand, some viruses of different families or even different groups used the same receptor (Figure 1). For example, HIV-2 and EBOV, from the family of *Retroviridae* (RT group) and *Filoviridae* (ssRNA group) respectively, both took CD209 molecule (CD209) (marked with an asterisk in Figure 1) as the receptor. On average, each receptor was used by more than two viruses. More specifically, among 119 virus receptors, forty-four of them were used by more than one virus (Figure 1 and Figure S1C); twenty-one of them were used by viruses of more than one family and fifteen of them were used by viruses of more than one group (Figure 1).

### Structural, functional, evolutionary and tissue-specific expression characteristics of mammalian virus receptors

To understand how the virus selects receptors, we systematically analyzed the structural, functional, evolutionary and tissue-specific expression characteristics of the mammalian virus receptor.

#### 1) The mammalian virus receptor were structurally diverse

We firstly investigated the structural characteristics of mammalian virus receptor proteins. As expected, all the mammalian virus receptor protein belonged to the membrane protein which had at least one transmembrane alpha helix (Figure S2A). Twenty-four of them had more than five helixes, such as 5-hydroxytryptamine receptor 2A (HTR2A) and NPC intracellular cholesterol transporter 1 (NPC1). The receptor protein was mainly located in the cell membrane. Besides, more than one third (43/119) of them were also located in the cytoplasm, and thirteen of them were located in the nucleus.

Then, the protein domain composition of the mammalian virus receptor protein was analyzed. The mammalian virus receptor proteins contained a total of 336 domains based on the Pfam database, with each viral receptor protein containing more than two domains on average (Figure S2B). This was significantly more than that of human proteins or human membrane proteins (p-values < 0.001 in the Wilcoxon rank-sum test). Some viral receptor proteins may contain more than 10 domains, such as complement C3d receptor 2 (CR2) and low density lipoprotein receptor (LDLR). The protein domains of the mammalian virus receptor protein could be grouped into 77 families in the Pfam database, suggesting the structure diversity of the mammalian virus receptor protein. The most commonly observed Pfam families were Immunoglobulin V-set domain, Immunoglobulin C2-set domain, Integrin beta chain VWA domain, Integrin plexin domain, and so on (Figure S2C).

#### 2) The mammalian virus receptor had high level of N-glycosylation

Glycosylation of protein is widespread in the eukaryote cell. We next characterized the glycosylation level of the mammalian virus receptor. N-glycosylation is the most common type of glycosylation. We found that 93 of 119 mammalian virus receptors were N-glycosylated with an average of 0.94 glycosylation sites per 100 amino acids (Figure 2A). It increased to 0.97 glycosylation sites per 100 amino acids for the human viral receptor (Figure 2A), among which 62 were N-glycosylated. Twelve human viral receptors were observed to have ten or more N-glycosylation sites, such as complement C3b/C4b receptor 1 (CR1) and lysosomal associated membrane protein 1 (LAMP1). Figure 2B displayed the modeled 3D-structure of HTR2A, the receptor for JC polyomavirus (JCPyV). Five N-glycosylation sites were highlighted in red on the structure, which were reported to be important for viral infection ^[30]^. For comparison, we also characterized the N-glycosylation level for the human cell membrane protein, human membrane proteins and all human proteins (Figure 2A). It was found they had a significantly lower level of N-glycosylation than that of human and mammalian virus receptors (p-values < 0.001 in the Wilcoxon rank-sum test), which suggests the importance of N-glycosylation for the viral receptor.

**Figure 2.**
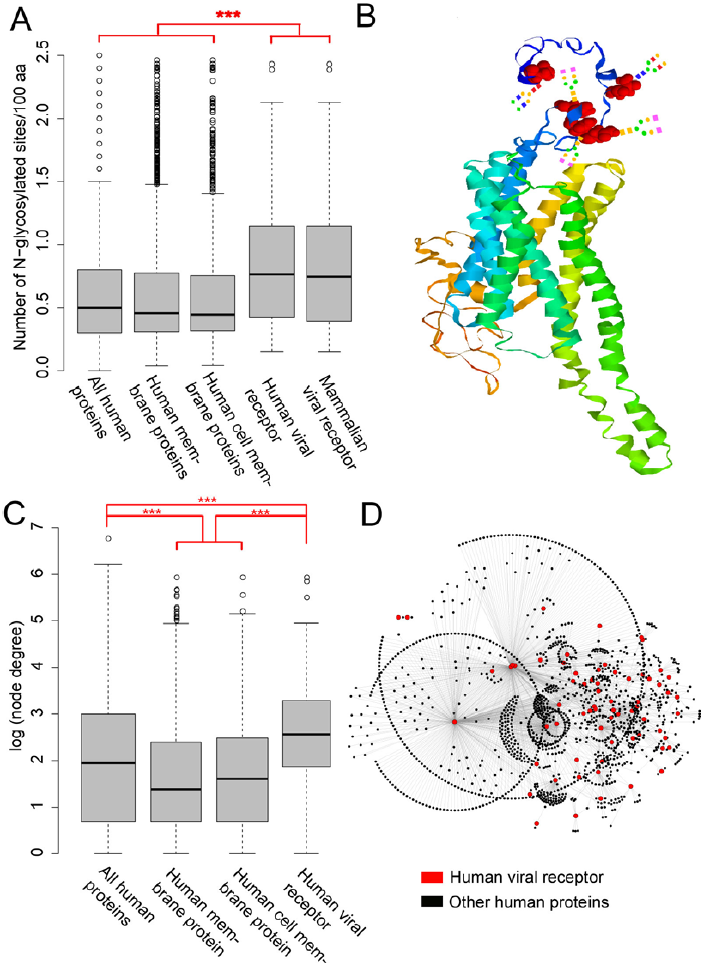
Analysis of N-glycosylation and protein-protein interactions of mammalian virus receptors. (A) Comparison of the N-glycosylation level between mammalian viral receptors, human viral receptors, human cell membrane proteins, human membrane proteins and all human proteins. For clarity, the outliers greater than 2.5 were removed. “***”, p-value < 0.001. (B) The modeled 3D-structure of HTR2A. Five N-glycosylation sites were highlighted in red. Artificial glycans were manually added onto the site. (C) Comparison of the degree of proteins between human viral receptors, human cell membrane proteins, human membrane proteins and all human proteins in the human PPIN. For clarity, the node degree was logarithmically transformed. “***”, p-value < 0.001. (D) Partial human PPI network composing of the PPIs which involved at least one viral receptor protein (colored in red).

O-glycosylation is also a common type of glycosylation. We found there was only a small fraction of mammalian virus receptors (14/119) with O-glycosylation. Besides, no significant difference was observed between the O-glycosylation level of mammalian virus receptor proteins and that of human proteins (Figure S2D).

#### 3) Functional enrichment analysis of the human virus receptor

We next attempted to identify the gene functions and pathways enriched in the mammalian virus receptor. As was mentioned above, 74 of 119 mammalian virus receptors belonged to the human. Besides, analysis showed that 36 of the remaining non-human mammalian virus receptors were homologs of the human virus receptor (Table S2). Therefore, we conducted the function enrichment analysis only for the human virus receptor based on the databases of Gene Ontology (GO) and KEGG. For the GO Cellular Component (Table S3), the human virus receptor was mainly enriched in the membranes and junctions, the latter of which included the adherens junction, cell-substrate junction, focal adhesion, and so on. For the GO Biological Process (Table S3), the human virus receptor was mainly enriched in the process of entry into the host. Besides, some terms related to the immune response were also enriched, such as “Regulation of leukocyte activation” and “Lymphocyte activation”. For the GO Molecular Function (Table S3), besides for the enrichment of terms related to the virus receptor activity, the human virus receptor was also enriched in terms of binding to integrin, glycoprotein, cytokine, and so on.

Consistent with the enrichment analysis of GO Cellular Component, the KEGG pathways of “Cell adhesion molecules”, “Focal adhesion” and “ECM-receptor interaction” were also enriched. Besides, the pathway of “Phagosome” was enriched (Table S3), which may be associated with viral entry into the host cell. Interestingly, some pathways associated with heart diseases were enriched, including “Dilated cardiomyopathy”, “Hypertrophic cardiomyopathy”, “Arrhythmogenic right ventricular cardiomyopathy” and “Viral myocarditis”.

#### 4) Human virus receptors had more interaction partners than other proteins

We next analyzed the protein-protein interactions (PPIs) which the mammalian virus receptor protein took part in. As the reason mentioned above, we only used the human virus receptor for PPI analysis. A human PPI network (PPIN) was constructed based on the work of Menche et al ^[31]^. It included a total of 13,460 human proteins that are interconnected by 141,296 interactions. The degree and betweenness of each protein in the PPIN were calculated, which could measure the importance of a protein in the PPIN. It was found that the degrees for human membrane proteins and cell membrane proteins were significantly smaller than those of other human proteins (Figure 2C & Figure S3A, p-value < 0.001 in Wilcox rank-sum test) in the PPIN. Similar observations could be found for the node betweenness in the PPIN (Figure S3B&C). However, the human virus receptor protein, a subset of the human cell membrane protein, was found to have significantly larger degrees and higher betweenness than other human proteins in the PPIN (Figure 2C and Figure S3, p-value < 0.001 in Wilcox rank-sum test). The median degrees for human virus receptors was 13 (Figure 2C), which was nearly twice as much as that of all human proteins. Six viral receptors were observed to have degrees larger than 100 (Figure 2D), including epidermal growth factor receptor (EGFR), heat shock protein family A member 8 (HSPA8), PHB, RPSA, CD4 molecule (CD4) and integrin subunit beta 1 (ITGB1). Since the viral receptor (colored in red in Figure 2D) interacted with lots of other human proteins (colored in black in Figure 2D) in PPIN, we further investigated the functional enrichment of these proteins by GO enrichment analysis. Interestingly, six of top ten enriched terms in the domain of Biological Process were related to protein targeting or localization (Table S3).

When looking at the interactions between viral receptors, we found that 38 of 74 viral receptors interacted with themselves. This ratio (38/74 = 51%) was much higher than that of human proteins (22%), membrane proteins (11%) and human cell membrane proteins (14%). However, we found the viral receptor tended not to interact with each other (Figure S3D). Among 74 human virus receptor proteins, 36 of them had no interactions with any other human virus receptor. There were only 50 PPIs between different human virus receptor proteins, with each viral receptor protein interacting with an average of only one other viral receptor protein.

#### 5) The mammalian virus receptor was not more conserved than other genes

Large degree of the human viral receptor in the human PPIN suggests the importance of them in cellular activity. Analysis showed that 11 human viral receptors belonged to the housekeeping gene. This ratio (0.15 = 11/74) was a little lower than that of housekeeping genes in all human genes (0.19 = 3804/20243), suggesting that the human viral receptor was not enriched in the housekeeping gene.

Then, we investigated the evolutionary conservation of mammalian virus receptors in 108 mammal species which were richly annotated in the NCBI Reference Sequences (RefSeq) database (see Methods). Over half of mammalian virus receptors had homologs (see Methods) in more than 100 mammal species (Figure S4A). We further calculated the pairwise sequence identities between the viral receptor and their homologs in mammal species. For nearly half of viral receptors, the average of pairwise sequence identities was higher than 0.8 (Figure S4B). For comparison, we also analyzed the evolutionary conservation of human proteins by randomly selecting 1000 human proteins from the NCBI RefSeq database. They were observed to have similar conservation level with that of human viral receptors (Figure S4C&D and Table S4).

#### 6) Viral receptors expressed higher than other proteins in 32 major human tissues

Since the virus has to compete with other proteins for binding to the receptor, proteins with high expression should be preferred by viruses as receptors. Thus we measured the average expression level of human viral receptors in 32 major human tissues (Figure 3). For comparison, those of human membrane genes, human cell membrane genes and all human genes were also displayed in Figure 3. As was shown clearly, the expression level of human cell membrane genes (in cyan) was similar to that of all human genes (in black) in these tissues. Both of them were lower than that of human membrane genes (in blue). However, the human viral receptor (in red), which was part of the human cell membrane gene, expressed much higher than other sets of genes in nearly all these tissues. On average, they had an expression level of 24 transcripts per million (TPM) in these tissues, while this was 8, 4 and 4 TPM for the human membrane gene, the human cell membrane gene and all human genes (see the black arrow in Figure 3).

**Figure 3.**
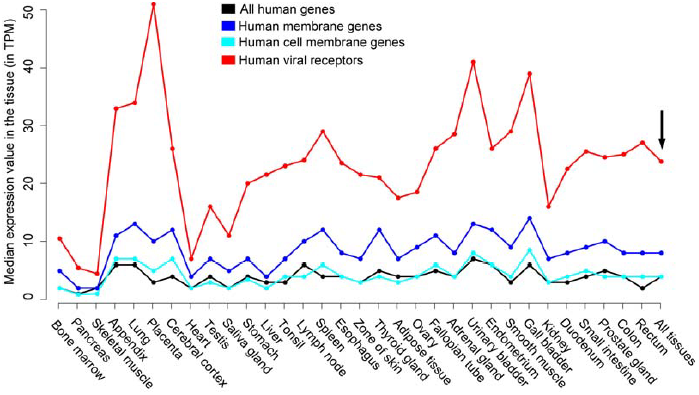
The average expression level of human viral receptors (red), human cell membrane genes (cyan), human membrane genes (blue) and all human genes (black) in 32 major human tissues. The expression level was measured with transcripts per million (TPM). The black arrow refers to the average expression level of genes in all 32 tissues.

### Viral receptors used by the same virus tended to co-evolve

The results mentioned above shows that a total of 51 viruses used more than one viral receptor. These viral receptors may work together or independently. We then analyzed the relationship between them. Structural analysis shows that except for integrins which generally work in heterodimer, few of viral receptors used by the same virus shared the same protein domain (data not shown). This suggests that when the virus expands the use of receptors, it tends to select structurally diverse proteins. We continued to analyze the co-evolution between mammalian viral receptors in 108 mammal species. We found that the average of Spearman Correlation Coefficients (SCCs, a measure of the extent of co-evolution) between viral receptors employed by the same virus was 0.54, which was significantly larger than that between other viral receptors (Figure 4A, p-value < 0.001 in the Wilcoxon rank-sum test). For example, SARS-CoV used four receptors, i.e., ACE2, CD209, C-type lectin domain family 4 member G (CLEC4G) and member M (CLEC4M). The average of SCCs between these four receptors was as high as 0.86. In addition, we analyzed the extent of co-expression between human viral receptors in 32 tissues. It was found that the extent of co-expression between viral receptors employed by the same virus was a little larger than that between other viral receptors, yet this difference was not statistically significant (Figure 4B, p-value > 0.1 in the Wilcoxon rank-sum test).

**Figure 4.**
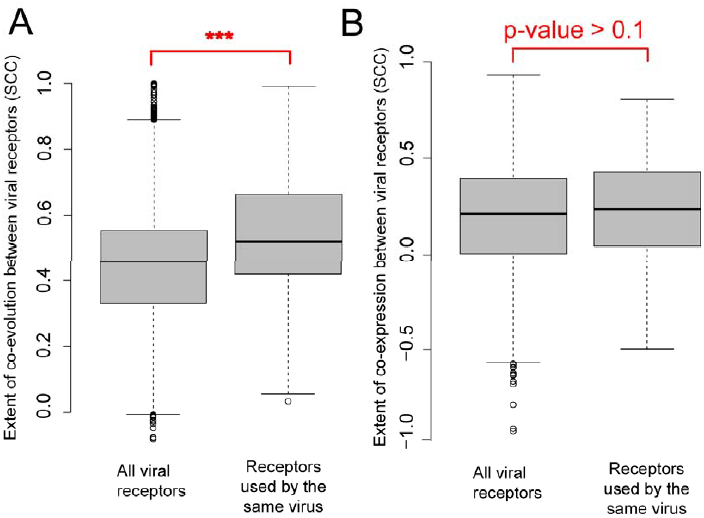
The co-evolution and co-expression of viral receptors. (A) Comparing the extent of co-evolution between mammalian virus receptors in 108 mammal species in the set of receptors used by the same virus and all mammalian virus receptors. “***”, p-value < 0.001 in the Wilcoxon rank-sum test. (B) Comparing the extent of co-expression between human viral receptors in 32 human tissues in the set of receptors used by the same virus and all human virus receptors.

### Analysis of the association between the tissue and host specificity of the virus and the viral receptor

Although there were plenty of studies about the tissue and host specificity of the virus and the viral receptor, there was still a lack of systematic analysis towards the association between them. Besides, few studies quantify such associations. Therefore, we further investigated systematically the association between the tissue and host specificity of the virus and the viral receptor.

#### 1) Viral receptor expressed higher in tissues infected by viruses than in those not infected

To investigate the association between the tissue specificity of the virus and tissue-specific expression of viral receptors, we manually compiled the tissue tropism of viruses from the literature or Wikipedia and obtained that in 32 human tissues for a total of 52 viruses (Table S5). Some viral receptors had high expression levels in most tissues, most of which were housekeeping genes, such as CD81 molecule (CD81) and ITGB1. While for most viral receptors, their expression levels varied much in different tissues. Analysis of the association between the tissue-specific expression of viral receptors and viral tissue tropism showed that the viral receptor expressed higher in the tissues infected by viruses (marked with asterisks in Table S5) than in those not infected, yet this difference was not statistically significant (p-value > 0.1 in the Wilcoxon rank-sum test) (Figure S5). For example, the neural cell adhesion molecule 1 (NCAM1), which was employed by the Rabies lyssavirus (RabiesV) as the receptor, expressed much higher in the tissue of Cerebral cortex (infected by RabiesV) than in other tissues not infected by the virus (Table S5).

#### 2) The viral receptor was a significant predictor in predicting viral cross-species in mammal species

Since the viral receptor determines the host specificity of the virus to a large extent, it is expected that the closer between the viral receptor and its homolog in a species, the more likely the virus which used the receptor would infect the species. To validate this hypothesis, we firstly calculated the sequence identities between the viral receptor and their homologs in 108 mammal species (Figure 5 and Table S6). For clarity, only 26 mammal species, which were frequently observed, were presented in Figure 5. Then, we compared the sequence identities between viral receptor proteins and their homologs in the species infected by the virus which used the receptor (marked with asterisks and triangles), and in those not infected by the virus. As expected, the former was significantly higher than the latter (Figure 6A, p-value < 0.001 in the Wilcoxon rank-sum test).

**Figure 5.**
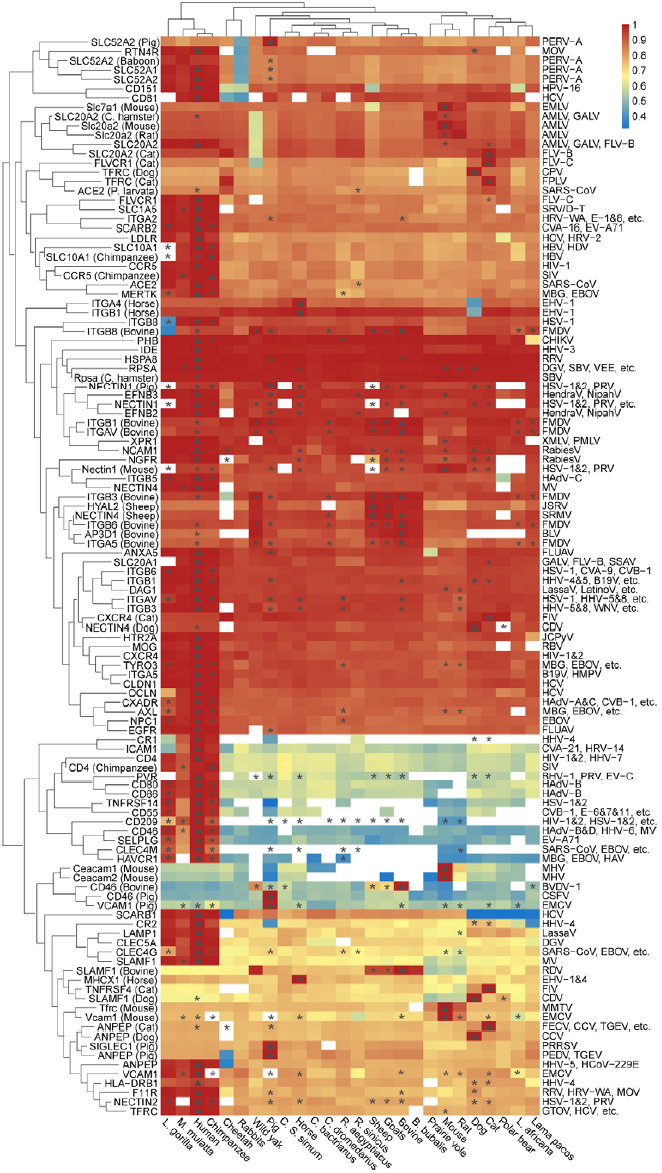
Analysis of the association between the host specificity of the virus and the viral receptor. It listed the sequence identities (colored according to the legend) between the viral receptor and its homologs in 26 mammal species (at the bottom). The white referred to no homologs in the species. The viral receptor and the virus which used them were displayed in the left and right side of the figure respectively. For the non-human mammalian virus receptor, the species it belongs to was presented in brackets. The asterisk referred to the viral infection of the mammal species based on Olival’s work, while the triangle referred to that based on our database. For more details, please see Table S6.

**Figure 6.**
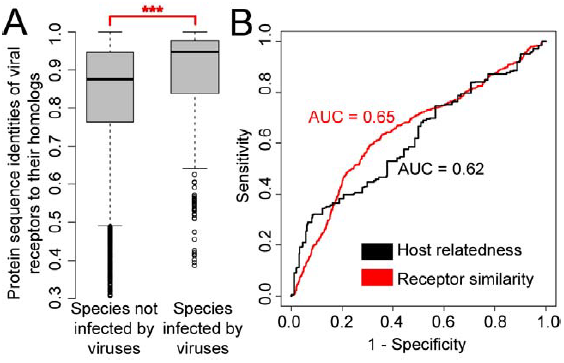
Quantify to what extent the viral receptor determine the host specificity of the virus. (A) Sequence identities between viral receptors and their homologs in mammal species infected by viruses and those not infected. “***”, p-value < 0.001. (B) The Receiver Operating Characteristic (ROC) curve for models of predicting viral cross-species in mammal species based on host relatedness (in black) and receptor sequence identity (in red). AUC, Area Under the ROC Curve.

Previous work by Olival et al. showed that phylogenetic relatedness of host was a major factor which influenced the cross-species of mammalian viruses ^[7]^. We compared the ability of the host relatedness and the viral receptor similarity in predicting viral cross-species in mammal species. The models based on the latter achieved an Area Under the ROC Curve (AUC) of 0.65 (Figure 6B), a little higher than that of the model based on the former (0.62), although this difference was not statistically significant (p-value = 0.13). This suggests that the viral receptor similarity should be a significant predictor in predicting viral cross-species in mammal species as well as the host relatedness.

Based on the results mentioned above, we continued to evaluate the risk of viral cross-species transmission in 108 mammal species based on viral receptors. Figure 5 shows that more than 40% of viral receptors, such as insulin degrading enzyme (IDE) and PHB, had high sequence identities with their homologs in most mammal species, suggesting that the virus using them may have a high probability to infect these species. On the other hand, some mammal species had homologs which were highly similar to most viral receptors, such as the primates (Chimpanzee, Macaca mulatta and Lowland gorilla). They may have a high risk of infection by the virus which used these receptors.

## Discussion

The viral receptor is essential for viral infection. By collecting the largest dataset ever reported about the mammalian virus-host receptor interactions, we systematically analyzed the structural, functional, evolutionary and tissue-specific expression characteristics of mammalian virus receptors. We found that the viral receptors were a subset of structurally and functionally diverse cell membrane proteins. They were enriched in GO terms and KEGG pathways related to junctions, adhesion and binding, which were typical features of viral receptors reported by previous studies ^[10, 12, 14, 15, 21]^. Besides, our analysis identified several novel features of the viral receptor. Firstly, the viral receptor had a higher level of N-glycosylation than other proteins. Then, what’s the relationship between glycosylation and viral receptor selection? As we know, glycosylation of proteins is widely observed in eukaryote cells ^[32]^. It plays an important role in multiple cellular activities, such as folding and stability of glycoprotein, immune response, cell-cell adhesion, and so on. Glycans are abundant on host cell surfaces. They were probably the primordial and fallback receptors for the virus ^[11]^. To use glycans as their receptors, a large number of viruses have stolen a host galectin and employed it as a viral lectin ^[11, 33]^. For example, the SJR fold, which was mainly responsible for glycan recognition and binding in cellular proteins, was observed in viral capsid proteins of over one fourth of viruses ^[33]^. Thus, during the process of searching for protein receptors, the protein with high level of glycosylation could provide a basal attachment ability for the virus, and should be the preferred receptor for the virus.

Secondly, our analysis showed that the viral receptor protein had a tendency to interact with itself and had far more interaction partners than other membrane proteins. Besides the function of viral receptor, the receptor protein functions in the host cell by interacting with other proteins of the host, such as signal molecules and ligands. Therefore, the virus has to compete with these proteins for binding to the receptor ^[15]^. The protein with less interaction partners are expected to be preferred by the virus. Why did the virus select the proteins with multiple interaction partners as receptors? One possible reason is that the receptor proteins are closely related to the “door” of the cell, so that many proteins have to interact with them for in-and-out of the cell. This could be partly validated by the observation that for the interaction partners of human viral receptors, six of top ten enriched terms in the domain of GO Biological Process were related to protein targeting or localization (Table S3). For entry into the cell, the virus also selects these proteins as receptors. Another possible reason is that viral entry into the cell needs cooperation of multiple proteins which were not identified as viral receptors yet. Besides, previous studies show that the virus could structurally mimic native host ligands ^[34]^, which help them bind to the host receptor. Thus, membrane proteins with multiple interaction partners have a larger probability to be used by viruses as receptors than other proteins.

Thirdly, the viral receptor was observed to have a much higher level of expression than other genes in each of the 32 human tissues. This may be directly related to the above finding that the viral receptor generally had multiple interaction partners: on the one hand, the viral receptor needs multiple copies to interact with multiple proteins; on the other hand, since the virus has to compete with other proteins for binding to the receptor, high expression of the receptor will facilitate the virus’s binding to the viral receptor.

The virus-receptor interaction is a major determinant of viral host range and tissue tropism. Previous case studies showed that the viral receptor expressed highly in the tissues infected by the virus ^[35, 36]^. Consistent with these studies, our systematic analysis found that the tissues with high expression of the viral receptor, and the mammal species with homologs highly similar to the viral receptor, were more possibly to be infected by the virus. However, the opposites were also observed. Some mammal species (or tissues) which had no receptor homolog (or low expression of the viral receptor) were also infected by the virus. These viruses may use other receptors not identified yet. Some mammal species (or tissues) with homologs highly similar to viral receptors (or high expression of the viral receptor) were observed to be not infected by the virus. This may be partly explained by the missed virus-host interactions in our data. Besides, it may also suggest that the host or tissue susceptibility to the virus is not solely determined by the viral receptor. More factors such as the host or tissue accessibility ^[7, 23]^, the cell defense system ^[37, 38]^ and the complex interaction between viral and host proteins ^[39, 40]^ may also influence viral infections.

There were some limitations within the study. Firstly, the viral receptor was biased towards the human, due to the bias of studies towards human viruses. Fortunately, the viral receptor was conserved in mammal species to a large extent, which may reduce the influence of this bias on the diversity of viral receptors. Secondly, the virus-host interactions were not complete due to limited surveys ^[7]^. According to the risk analysis of viral cross-species based on viral receptors, much more mammal species may be infected by the mammalian virus analyzed in this study. High attention should be paid to this risk. Thirdly, due to the difficulties of identifying viral receptors ^[17, 41, 42]^ the database of mammalian virus-host receptor interaction was still limited in its size, which hindered us from a more comprehensive survey of the correlation characteristics between viruses and viral receptors. More effective methods, either experimental or computational ^[34]^, should be developed for identifying viral receptors, while the characteristics identified in this study may help such endeavors.

Overall, the structural, functional, evolutionary and tissue-specific expression characteristics identified here should not only deepen our understanding of the viral receptor selection, but also help for development of more effective methods for identifying viral receptors. Besides, evaluating the risk of viral cross-species infection based on the viral receptor could also help for early warning and prediction of viral zoonotic diseases.

## Materials and Methods

### Database of mammalian virus-host receptor interaction

The data of mammalian virus-host receptor interaction were compiled from three sources: firstly, the literature related to viral receptors (a total of 1303 papers) were downloaded from NCBI Pubmed database ^[43]^ by searching “virus receptor” [TIAB] or “viral receptor”[TIAB] on August 14^th^, 2017. The mammalian virus and their related host receptors were manually extracted from the literature; secondly, part of viral receptors were directly obtained from the database of ViralZone ^[44]^ on September 9^th^, 2017; thirdly, proteins annotated with one of GO terms “virus receptor activity”, “viral entry into host cell” and “receptor activity” in UniprotKB database ^[45]^ were collected on August 14^th^, 2017, and manually checked later. In combination, a database was created with 268 pairs of mammalian virus-host receptor interaction, which included 128 unique viral species or sub-species and 119 viral receptors (Table S1).

### Analysis of structural features of viral receptors

The number of transmembrane alpha helix of the mammalian virus receptor was derived from the database of UniprotKB and the web server TMpred ^[46]^. The location for the viral receptor was inferred from the description of “Subcellular location” for the receptor protein provided by UniProtKB, or from the GO annotations for them: the viral receptors annotated with GO terms which included the words of “cell surface” or “plasma membrane” were considered to be located in the cell membrane; those annotated with GO terms which included the words of “cytoplasm”, “cytosol” or “cytoplasmic vesicle”, or shown to be in the cytoplasm in UniProtKB, were considered to be located in the cytoplasm; those annotated with GO terms “nucleus” (GO:0005634) or “nucleoplasm” (GO:0005654) were considered to be located in the nuclear. The Pfam family, the N-glycosylation and O-glycosylation sites for the protein were obtained from the database of UniprotKB.

For comparison, the human proteins and their related structural characteristics were obtained from the database of UniProtKB/SwissProt on November 24^th^, 2017. The proteins which had at least one transmembrane alpha helix were considered as membrane proteins. The membrane proteins which were shown to be located in the cell membrane were considered as cell membrane proteins. In total, we obtained 20243 human proteins, 5187 human membrane proteins and 2208 human cell membrane proteins.

The 3D structure for the viral receptor HTR2A were modeled with the help of I-TASSER ^[47]^ based on the protein sequence of HTR2A (accession number in the database of UniProt: P28223). The best model was selected, and visualized in RasMol (version 2.7.5) ^[48]^.

### Functional enrichment analysis

The GO function and KEGG pathway enrichment analysis for the human viral receptor were conducted with functions of *enrichGO()* and *enrichKEGG()* in the package “clusterProfiler” (version 3.4.4) ^[49]^ in R (version 3.4.2) ^[50]^.

### Protein-protein interaction (PPI) network analysis

The human PPI network (PPIN) was constructed based on the work of Menche et al [31]. The degree and betweenness for each protein in the PPIN were calculated with functions of *degree()* and *betweenness()* in the package “igraph” (version 1.0.0) ^[51]^ in R (version 3.2.5). The network was displayed with Cytoscape (version 2.6.2) ^[52]^.

For robustness of the results, we also conducted PPI analysis based on the human PPIs derived from the database of STRING ^[53]^ on November 7, 2017. The human PPIN was built based on the PPIs with median confidence (combined score equal to or greater than 0.4). It included 710,188 PPIs and 17,487 proteins which could be mapped to NCBI gene ids. Similar to those mentioned above, the viral receptor protein was observed to have far more interaction partners and higher betweenness than other proteins in the human PPIN (Figure S3A&C).

### Evolutionary analysis

To identify the homolog of the mammalian virus receptor in other mammal species, the protein sequence of each viral receptor was searched against the database of mammalian protein sequences, which were downloaded from NCBI RefSeq database on ^[54]^October 10^th^, 2017, with the help of BLAST (version 2.6.0) ^[55]^. Analysis showed that in the database of mammalian protein sequences, there were 108 mammal species which were richly annotated and had far more protein sequences than other mammal species (Table S7). Therefore, only these 108 mammal species were considered in the evolutionary analysis. Based on the results of BLAST, the homolog for the viral receptor was defined as the hit with E-value small than 1E-10, coverage equal to or greater than 80% and sequence identity equal to or greater than 30%. Only the closest homolog, i.e., the best hit, in each mammal species was used for further analysis. For measuring the conservation level of viral receptors, two indicators were used. The first indicator was the number of mammal species with homolog of the viral receptor in 108 mammal species. The other indicator was the average of the pairwise sequence identities between the viral receptor and its homologs in 108 mammal species. For comparison, 1000 human protein sequences were randomly selected from the NCBI RefSeq database (Table S4). Similar methods as above were utilized to calculate the indicators of conservation level for these proteins.

For analysis of co-evolution between viral receptors, firstly for each viral receptor, a phylogenetic tree was built based on the protein sequences of the receptor and its homologs in 108 mammal species with the help of phylip (version 3.68) ^[56]^. The neighbor-joining method was used with the default parameter. Then, the genetic distances between the viral receptor and their homologs were extracted from the phylogenetic tree with a perl script. Finally, for a pair of viral receptors, the spearman correlation coefficient (SCC) was calculated between the pairwise genetic distances of viral receptors and their homologs, which was used to measure the extent of co-evolution between this pair of viral receptors.

The set of housekeeping gene in human was adapted from the work of Eisenberg et al^[57]^ A total of 3804 genes were identified as the housekeeping gene.

### Analysis of the tissue-specific expression of human viral receptors

The expression level for human viral receptors and other human genes in 32 human tissues were derived from the database of Expression Atlas ^[58]^. For analysis of the association between viral infection and tissue-specific expression of viral receptors, we manually compiled the tissue tropism of viruses from the literature or Wikipedia and obtained that in 32 human tissues for a total of 52 viruses which used a total of 46 receptors (Table S5). When comparing the expression level of human viral receptors and other set of human genes in 32 human tissues, to reduce the influence of extreme values, the median instead of the mean of the expression values was used to measure the average expression value of a gene set in a tissue.

The SCCs between the expression values of viral receptors in 32 tissues were calculated to measure the extent of co-expression between viral receptors.

### Analysis of the host specificity of the virus and the viral receptor

The mammalian virus-host interactions were primarily adapted from Olival’s work ^[7]^. One hundred and fifteen viruses in our database and 61 of 108 richly annotated mammal species could be mapped to those in Olival’s work (Table S8). These 115 viruses used a total of 116 viral receptors. The sequence identities of these viral receptor proteins to their related homologs in the corresponding mammal species were presented in Table S6.

For comparison, we also extracted genetic distances (host relatedness) between the mammal species and the viral host with reported receptors based on Olival’s work (Table S9). Then, the genetic distance of the mammal species to the viral host with reported receptors, and the sequence identity of the receptor homolog in the mammal species to the viral receptor protein, was respectively used to predict whether a mammal species could be infected by the virus which infected the host with reported receptors. The method of Receiver Operating Characteristic (ROC) curve was used to evaluate and compare their performance with the functions of *roc(), auc(), roc.test()* and *plot.roc()* in the package of “pROC” ^[59]^ in R (version 3.2.5).

### Statistical analysis

All the statistical analysis was conducted in R (version 3.2.5) ^[50]^. The wilcoxon rank-sum test was conducted with the function of *wilcox.test()*.

## Acknowledgements

This work was supported by the National Key Plan for Scientific Research and Development of China (2016YFD0500300 and 2016YFC1200200), the National Natural Science Foundation of China (31500126, 31671371, 81730064 and 81571985), National Science and Technology Major Project (2017ZX10202201) and the Chinese Academy of Medical Sciences (2016-I2M-1-005). The authors would like to thank Pro. Xiangjun Du in Sun Yat-sen University for helpful suggestions.

The authors have declared that no competing interests exist.

## Author contributions

HZ, TJ and YP conceived and designed the study. ZZ and ZZZ did the computational analysis. WC, ZC, ZT and BX compiled the database of mammalian viruses and their related receptors, and the tissue tropism of viruses. AW and XYG directed the computational analysis about the tissue and host specificity of viruses. YP and ZZ wrote the paper. AW, XYG, XHG, ZT, ZX, TJ and HZ reviewed and edited the manuscript. All authors read and approved the manuscript.

